# Identification of *Amblyomma americanum* antigens after vaccination with tick extracellular vesicles in white-tailed deer

**DOI:** 10.1101/2025.02.03.636374

**Authors:** Adela Oliva Chávez, Julia Gonzalez, Cristina Harvey, Cárita de Souza Ribeiro-Silva, Brenda Leal-Galvan, Kelly A. Persinger, Sarah Durski, Pia U. Olafson, Tammi L. Johnson

## Abstract

Anti-tick vaccines represent a promising alternative to chemical acaricides for the management of ticks on wildlife; however little progress has been made to produce a vaccine effective in wild hosts that are critical for tick reproduction, such as the white-tailed deer (*Odocoileus virginianus*; WTD). To date, most tick antigens have been tested using laboratory models (i.e. rabbits); however, their expression in wild hosts has not been confirmed. We recently tested *Amblyomma americanum* salivary (SG) and midgut (MG) extracellular vesicles (EVs) as vaccine candidates in WTD, which resulted in on-host female tick mortality. Using a proteomic approach, we show that these SG- and MG-EVs contain a “core-cargo” enriched in chaperones, small GTPases, actin and actin-related proteins, and other proteins previously reported in small EVs. Label-free quantitative proteomics showed significant differences in protein cargo between MG and SG-EVs (333 proteins out of 516). Pre-vaccinated and day 57 post-injection serum samples from three vaccinated and one control WTD were used to immuno-precipitate antigenic proteins from SG- and MG-EV preparations. Proteomic analysis of immunoprecipitated proteins identified thirty antigens with potential for use in anti-tick vaccines, seven of which we have categorized as high priority. These proteins represent promising candidates for anti-tick vaccine design in WTD and other wildlife hosts.

## 1. Introduction

Ticks are important vectors of human and animal pathogens. In recent years, there has been a rise in the incidence of tick-borne diseases in the US [1]. Further, milder winters due to climate change will likely affect the geographic range of ticks, and consequently, tick-borne diseases [2]. Among the tick species that most frequently bite humans in the US, *Amblyomma americanum* Linnaeus (Acari: Ixodidae; lone star tick) is known to transmit bacteria and viruses, and its bite may cause a delayed hypersensitivity to red meat in humans [3]. Additionally, this tick affects domestic animals, causing important economic damage and mortalities. For example, *A. americanum* infestations of cattle can result in weight loss in the absence of acaricide treatment [4]. This tick species is an aggressive biter that is expanding into the upper Midwest and Northeast US [5] and it is predicted that increasing temperatures may permit expansion as far north as south Quebec, Canada [6,7]. Thus, management tools that will allow the control of this tick species on wildlife and domestic animals are desirable.

White-tailed deer (*Odocoileus virginianus* Zimmermann; WTD) are considered the most suitable host for *A. americanum* [8] and their presence within a landscape positively correlates with the abundance of several tick species, including *Ixodes ricinus* Linnaeus (Acari: Ixodidae) [9], *Ixodes scapularis* Say (Acari: Ixodidae) [10], and *A. americanum* [11]. Deer densities also correlate with the incidence of *A. americanum-*borne diseases [12] and they serve as reservoirs for the human pathogens *Ehrlichia chaffeensis,* the causative agent of Human Monocytic Ehrlichiosis [13], and *E. ewingii* [14], a bacterium that infects neutrophils also causing Human Ehrlichiosis [15]. Given the importance of WTD in the density of several tick species and the transmission of tick-borne pathogens, interest in the development of management tools, including anti-tick vaccines, has grown [16]. Current studies treating WTD with permethrin using 4-poster devices showed a significant decrease in host-seeking larvae, nymphs, and adults. Moreover, these treatments resulted in a reduction in *Ehrlichia* spp. infected *A. americanum* [17]. However, 4-poster devices present several complications, including their limited effectiveness at a broader geographic scale, difficulties with regulations in their proximity to residential areas, high operational costs, the banning of baiting in several states throughout of the US due to chronic wasting disease (CWD), withdraw periods due to meat consumption, and limited adoption by the public [18]. According to a study done in Connecticut and New York, only 37% of residents would support the placement of 4-post devices within their properties, mostly due to safety concerns [19], thus, hindering their implementation. Although resistance to acaricides has not been reported in *A. americanum* populations, reports demonstrated potential development of tolerance against permethrins in wild *A. americanum* populations recovered from farmed and wild WTD [20]. Therefore, including immunological approaches (i.e. anti-tick vaccines) into integrated tick management programs is desirable [21]. Nevertheless, there are no current vaccines targeting *A. americanum* on WTD.

We recently tested the efficacy of tick salivary (SG) and midgut (MG) extracellular vesicles (EVs) as vaccine candidates in WTD, showing strong serum conversion [22]. Extracellular vesicles are small lipid-rich blebs that are secreted within tick saliva and hemolymph [23–25]. These vesicles are essential for tick feeding, pathogen transmission, and pathogen acquisition [24,26]. Interestingly, vaccinations with SG- and MG-EVs resulted in long-lasting circulating antibodies that were detected as far as 13 months after the last booster injection and led to 50.3% control of *A. americanum* females [22]. However, the antigens that elicited these immune responses remain to be identified. The objective of this study was to characterize the proteins that elicited an immune response within the vaccinated animals in our previous experiments.

## 2. Materials and Methods

### 2.1. Label-free quantitative proteomic analysis

Salivary (SG) and midgut extracellular vesicles (used to vaccinate WTD were isolated from *ex vivo* organ cultures from *A. americanum* females fed for 5 days on WTD as previously described [22]. Three aliquots of 365 µl of SG- or MG-EVs resuspended in 1X Phosphate Buffered Saline (PBS) were submitted to the Gehrke Proteomics Center at the University of Missouri for label-free quantitative proteomic analysis. Samples were processed as follows:

#### 2.1.1. Sample preparation

Protein pellets were washed with 80% acetone in water and centrifuged at 16,000 x g. Pellets were suspended in 10 µl urea buffer (6M urea, 2M thiourea, 100mM ammonium bicarbonate, pH 8.0). Protein was digested at 37 °C overnight with 0.3 µg trypsin. A second digestion was performed the next day with 0.2 µg Trypsin for 3 hours. Peptides were desalted using 100 µl C18 tips (Pierce). Lyophilized peptides were resuspended in 10 µl 5% acetonitrile and 0.1% formic acid. Peptide concentration was determined using a Pierce Quantitative Colorimetric Peptide assay (Thermo Fisher Scientific) and an equal amount of peptides were transferred into autosampler vials. The auto-sampler was cooled to 7°C during the run.

#### 2.1.2. Mass spectrometry

Peptides were analyzed by mass spectrometry, using the following procedure: 1 µl injection was made onto a 20 cm long x 75 µm inner diameter pulled-needle fused-silica analytical column packed with Waters BEH-C18, 1.7 µm reversed phase resin. Peptides were separated and eluted from the analytical column with a gradient of acetonitrile at 300 ηl/min. Liquid chromatography (LC) gradient conditions were as follows: initial conditions were 3% B (A: 0.1% formic acid in water, B: 99.9% acetonitrile, 0.1% formic acid), followed by a 10 min gradient to 17% B. Then 17-25% B over 20 min, 25-37% B over 10 min, 37-80% B over 6 min, ending with 3 flushing cycles (80% B-40% B for 2 minutes) and finally back to initial conditions (3% B) for 2 min and held for 4 min to equilibrate the column. Total run time was 60 min. Data was acquired on a timsTOF-PRO mass spectrometer (Bruker).

MS data were collected in positive-ion data-dependent (DDA) Parallel Accumulation-Serial Fragmentation (PASEF) mode over an m/z range of 100 to 1700. PASEF and TIMS were set to “on”. One MS and ten PASEF frames were acquired per cycle of 1.17 sec. Target MS intensity for MS was set at 10,000 with a minimum threshold of 2500. A charge-state-based rolling collision energy table from 20-59 eV was used. An active exclusion/reconsider precursor method with release after 0.4 min was used. If the precursor (within mass width error of 0.015 m/z) was > 4X signal intensity in subsequent scans, a second MS/MS spectrum was collected. Isolation width was set to 2 (<700m/z) or 3 (800-1500 m/z).

#### 2.1.3. Data analysis

The DDA-PASEF individual runs were analyzed with Fragpipe version 18.0 using the ‘LFQ-MBR’ workflow, which included the database search against Uniprot *Ixodes scapularis* (20,486 entities) with MSFragger version 3.5. This database enabled us to perform enrichment analyses based on the annotated *I. scapularis* genome. Although an annotated genome is available for *A. americanum* through NCBI (GCA_030143305.2), its proteins are not accessible through Panther [27]. Philosopher version 4.4.0 was used for peptide-spectrum match (PSM) validation and protein inference and IonQuant for MS1-level label-free quantification. For MS1-level quantification, the match between runs (MBR) and normalization were enabled along with the MaxLFQ algorithm, and a minimum number of 2 ions was required for quantifying a protein in IonQuant. For feature detection in IonQuant, 10 ppm, 0.4 min, and 0.05 1/k0 were set for m/z, retention time and ion mobility tolerances, respectively.

Data were searched with trypsin as the digestive enzyme, two missed cleavages were allowed, carbamidomethyl cysteine was included as a fixed modification, and oxidized methionine and N-terminal acetylation were used as variable modifications. The mass tolerances were set at 20 ppm for precursor ions and 0.1 Da for fragment ions. Data were exported and further analyzed with Perseus version 1.6.5.0. Only proteins with at least one unique peptide/protein and four total spectral of each whole set were used for quantitative comparison. Significant differences in quantification were evaluated by Student’s T-test followed by Benjamini Hochberg correction.

### 2.2. Extracellular vesicle cargo comparison and pathway enrichment analysis

Proteomic data were exported into Excel spreadsheets for analysis. Normalized intensity counts of the spectra (Supplementary File S1) were used to identify the cargo within each EV type. The UniProt ID from proteins with spectra counts in all three biological replicates and technical duplicates were filtered and the SG- and MG-EV cargo were compared using https://molbiotools.com/listcompare.php to identify shared and unique proteins.

UniProt IDs from shared proteins (Supplementary File S2) and proteins that showed differential quantitative enrichment by 1.5 fold (Supplementary File S3) were used to analyze the enrichment of the protein based on biological function, molecular function, and protein class, using Panther 19.0 (https://pantherdb.org; version released 06/20/2024 [27]) based on the available *I. scapularis* genome. Statistical significance was evaluated using Fisher exact test and corrected using a false discovery rate (FDR) of less than 0.05. Fold enrichment values were graphed with GraphPad (Prism).

### 2.3. Western blot analysis

Western blot analysis was used to confirm the presence of Hsp70, a protein enriched in small and large exosomes [28], and the absence or minimal presence of Calnexin, part of the endoplasmic reticulum but only present in some EV populations [29]. Proteins from EVs and organs were processed and separated by SDS-PAGE as we previously described [30]. Proteins were transferred onto Immun-Blot® polyvinylidene fluoride (PVDF) membrane sandwiches (Bio-Rad) and blocked overnight with 5% dry milk (Research Products International) in 1X PBS. Membranes were probed with rabbit anti-Hsp70 polyclonal antibodies (1:20,000; ProteinTech) or rabbit anti-Calnexin polyclonal antibodies (1:10,000; Millipore sigma) diluted in 10 mL 1X PBS + 25 µl 5% milk + 5 µl Tween 20 and detected with goat anti-rabbit polyclonal antibodies horse-radish peroxidase (HRP) labeled (1:10,000; Abcam) diluted in 10 mL 1X PBS + 25 µl 5% milk + 5 µl Tween 20. Bands were detected with Pierce^®^ ECL Western Blot Substrate (Thermo Fisher Scientific) and visualized in an Azure 300 Imager (Azure Biosystems).

### 2.4. Immunoprecipitation and label-free quantification of antigenic proteins

Serum samples from pre-vaccination and 7 days post-3rd booster (day 57 of the study) were obtained from three WTD vaccinated with SG- and MG-EVs (#924, #929, and #934) and one control WTD (#930) from our previous study [22]. These sera were submitted to the Gehrke Proteomics Center at the University of Missouri along with SG- and MG-EVs for immunoprecipitation and proteomic analysis of proteins bound by antibodies (antigenic). Identification of antigenic proteins with animal serum #929 and control (WTD #930) was performed on a different occasion from the immunoprecipitation with serum samples #924, #934, and control #930. The procedures for both immunoprecipitation experiments are described below:

#### 2.4.1. Immunoprecipitation

*Immunoprecipitation #929/930:* MG-EVs (60 µl) and SG-EVs (75 µl) were combined 1:1 with proteinA equilibration buffer before freeze/thawing three times, followed by sonication in a water bath for 5 min three times. Samples were aliquoted in triplicate (18 µl for MG-EVs and 20 µl for SG-EVs) and incubated with pre-vaccination serum (4 µl for WTD#929 serum or 8 µl for WTD#930 serum) at 4°C for 1 hr at mild agitation. Pierce™ Protein A/G Agarose (Thermo Fisher Scientific) was pre-equilibrated by washing three times in buffer. Fifty (50) µl of 50% Protein A/G Agarose was added into each sample and incubated for 30 min at 4 °C. Samples were centrifuged for 5 min at 2,000 x g and flow through was recovered. Protein A/G agarose-EV-pre-vaccination complexes were retained to identify potentially sticky proteins and unspecific identifications. The flow-through was incubated with post-vaccination serum (10 µl for WTD#929 serum or 9 µl for WTD#930 serum) at 4°C for 1 hr at mild agitation. Protein A/G Agarose was added (50 µl of 50%) and incubated for 30 min at 4 °C. Samples were centrifuged for 10 minutes at 16,000 x g. Precipitates were washed 3 times with equilibration buffer followed by 3 x washes with 1x PBS.

*Immunoprecipitation #924/934/930:* Immunoprecipitation was performed as detailed above, except that 95 µl of MG-EVs and SG-EVs were combined with proteinA equilibration buffer 1:1 and 30 µl were equally aliquoted in duplicates. An equal volume of pre-vaccination serum (10 µl) and post-vaccination serum (10 µl) was used for each animal (WTD #924, #934, and control WTD#930).

#### 2.4.2. Mass spectrometry

Immunoprecipitates were digested on beads with trypsin and peptides were desalted using C18 tips (Pierce). The eluates were lyophilized and resuspended in 10 µl of 5% acetonitrile and 0.1% formic acid.

*LC/MS:* Samples immunoprecipitated with serum from vaccinated WTD #929 and #930 were originally acquired using data-dependent acquisition (DDA). LCMS acquisition was performed as follows: PASEF and TIMS were set to “on”. One MS and ten PASEF frames were acquired per cycle of 1.17sec (∼1MS and 120 MS/MS). Target MS intensity for MS was set at 10,000 counts/sec with a minimum threshold of 250 counts/s. A charge-state-based rolling collision energy table was used from 76-123% of 42.0 eV. An active exclusion/reconsider precursor method with release after 0.4min was used. If the precursor (within mass width error of 0.015 m/z) was >4X signal intensity in subsequent scans, a second MSMS spectrum was collected. Isolation width was set to 2 (<700m/z) or 3 (800-1500 m/z). This allowed us to identify precursors to reduce error during Data-Independent-Acquisition (DIA).

Samples immunoprecipitated with serum from vaccinated WTD #929 were later analyzed using DIA settings, as follow: 1 µl of resuspension was loaded onto a pepsep25 column (25 cm x 75 µm x 1.9 µm ReproSil-AQC18; Bruker). Peptides were separated with a gradient of acetonitrile at 300 ηl/min with the following LC gradient conditions: initial conditions were 3% B (A: 0.1% formic acid in water, B: 99.9% acetonitrile, 0.1% formic acid), followed by 10 min gradient to 17% B. Then 17-25% B over 5 min, 25-37% B over 5 min, 37-80% B over 3 min, end with three flushing cycles (80%-40% B for 1 min) and finally back to initial conditions (3% B) for 1 min and hold for 2 min to equilibrate the column. The total run time was 30 min. MS data were collected using DIA-PASEF over an m/z range of 300.5 to 1349.5 and a mobility range of 0.65 to 1.45. Fifty DIA windows were collected (of varying m/z width) for a total cycle time of 2.76 sec. An ion-mobility-based collision energy was used with 20eV at 0.85 and 59eV at 1.30. This procedure using DIA settings was also used for samples immunoprecipitated with WTD #924, #934, and #930 but was completed on a different date.

#### 2.4.3. Data analysis

Data acquired during DDA LC/MS were submitted to the PEAKS-11 search engine for protein identifications. The NCBI-Ixodes database was searched with trypsin-specific as enzyme, allowing for 2 missed cleavages, and using carbamidomethyl-Cys as fixed modification and oxidized methionine as variable modification. The mass tolerance for precursor ions was set at 20 ppm and 0.1 Da on-fragment ions. A decoy database was automatically generated and searched for FDR calculation. Search results files were filtered for 0.1% FDR (peptide false discovery rate).

The raw data were copied to Spectronaut V18 (Biognosis Inc) server and searched against an NCBI-Ixodes FASTA database using the direct DIA approach incorporating the Pulsar algorithm. Spectronaut default parameters were used for the search with trypsin as enzyme, Carbamidomethyl-Cys as fixed modification, and Oxidized-Met and deamidated-NQ as variable modifications. Protein quantity was reported as MS2Quantity (sum of peptide fragments intensities from CID MS/MS matching each protein). Quantitative data from WTD #924/934/930 immunoprecipitations were statistically evaluated as follows: data (MS2Quantity, sum of peptide fragment abundances per protein) were imported into Perseus V16.0.1 and grouped into 12 groups (3 “pre” SG & MG; and 3 “vacc” MG & SG). Data were filtered for an MS2Quantitiy of ≥10 in two samples of at least one group and data sum normalized (protein abundance/total abundance) in each sample. T-tests were then conducted with two valid values in at least one group to compare pre- and post-vaccine for each animal and each tissue sample. The DIA-PASEF mass acquisition for the immuno-precipitates with WTD #929 serum resulted in high variation and could not be analyzed statistically.

### 2.5. Data availability

The mass spectrometry proteomics data associated with this manuscript have been deposited to the ProteomeXchange Consortium via PRIDE [31] repository with the dataset identifier PXD058874.

### 2.6. Antigens classification

Proteins identified during the immunoprecipitation were classified into three different groups: high, medium, and low priority, using the criteria described below.

*High priority:* Proteins were precipitated with at least two vaccinated WTD serum samples; proteins were identified in the samples precipitated with post-vaccinated serum and not present in pre-vaccination pool (unique to vaccinated); statistically significant by adjusted t-test analysis; and absent in precipitates using control serum (WTD #930).

*Medium priority*: Proteins were precipitated with at least two vaccinated WTD serum samples; proteins were ≥5-fold enriched in precipitates processed with post-vaccination serum when compared to pre-vaccination serum; statistically significant by adjusted t-test analysis; and absent in precipitates using control serum (WTD #930).

*Low priority*: Proteins were precipitated with at least two vaccinated WTD serum samples; proteins were identified in the samples precipitated with post-vaccinated serum and not present in pre-vaccination pool (unique to vaccinated); however, proteins presented variability in spectra intensity levels and were not significant by adjusted t-test analysis; and absent in precipitates using control serum (WTD #930).

The potential identity of proteins without known function (hypothetical or putative proteins) was investigated by Position specific iterated (PSI) Basic Local Alignment Search Tool (BLAST) [https://blast.ncbi.nlm.nih.gov/] using *I. scapularis* as the target species.

## 3. Results

### 3.1. Extracellular vesicles secreted by different organs share a core set of proteins

Recent studies demonstrated that certain proteins are associated with vesicle subtypes independent of the secreting cell origin [28]. These proteins can be used as markers to classify the different EV subpopulations and the International Society for Extracellular Vesicles (ISVE) has designated them for the validation of EVs’ identities [29]. Although our and other studies identified several of these markers in salivary and hemolymph EVs, using western blotting and proteomic analysis [23–25], it remained to be determined whether tick vesicles secreted by different organs shared a core proteomic cargo. Proteomic analysis of the SG- and MG-EVs used for our previous vaccination experiments [30] identified 1,539 proteins (Supplementary File S1). From these, 914 and 360 proteins were present in all biological and technical replicates of MG- and SG-EVs, respectively (Figure 1A). To determine the core protein cargo transported within *A. americanum* SG- and MG-EVs, we compared the proteins identified in all technical and biological replicates in each EV type. This core cargo consisted of 340 proteins that were shared between SG- and MG-EVs (Figure 1A). Enrichment analysis of the proteins found within the core cargo showed an over-representation of proteins involved in glycogen metabolic process (GO:0005977), barbed-end actin filament capping (GO:0051016), purine metabolic process (GO:0009167), and other metabolic processes (Figure S1A). Molecular functions that were enriched included constituents of ribosome (GO:0003735), translation elongation factor (GO:000374), ATP binding (GO:0005524), proteins disulfide isomerase activity (GO:0003756), and included GTPase activity (GO:0003924) (Figure S1B). Among the protein class enriched within both vesicles chaperonin (PC00073), Hsp90 family chaperone (PC00028), Hsp70 family chaperone (PC00027), actin and actin related protein (PC00039), ribosomal proteins (PC00202), vesicle coat proteins (PC00235), small GTPases (PC00208), and others were identified (Figure 1B). The presence of Hsp70 within both EVs samples was confirmed by western blot analysis, as well as the absence of the endoplasmic reticulum protein Calnexin (Figure 1C). This confirms that several markers used to classify EVs in other systems are shared among vesicles secreted by different organs in *A. americanum*.

**Figure 1.**
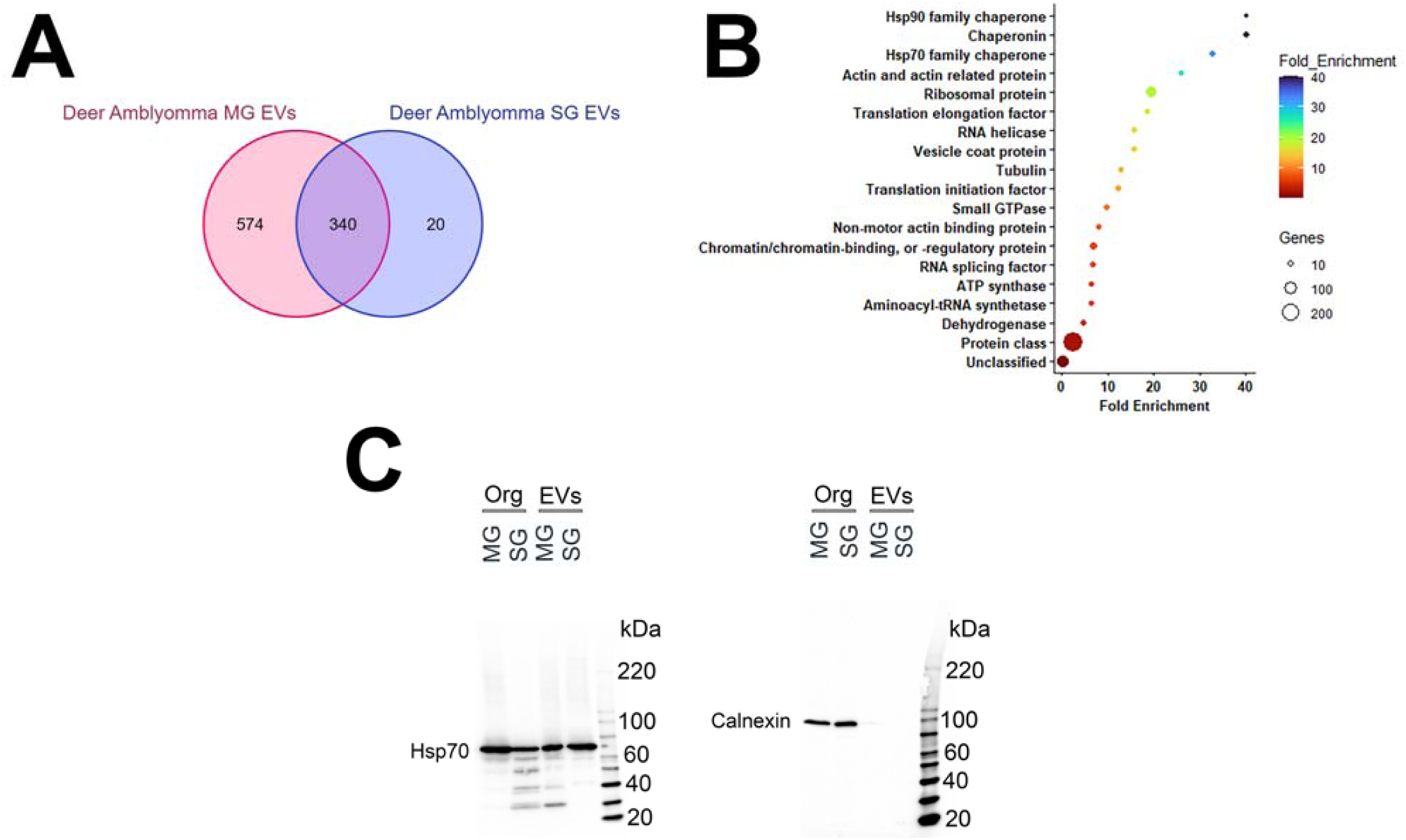
Shared proteomic cargo packed within salivary and midgut extracellular vesicles secreted by *Amblyomma americanum* females fed on white-tailed deer. Extracellular vesicles (EVs) from salivary glands (SG) and midguts (MG) were characterized by mass spectrometry using Parallel Accumulation-Serial Fragmentation (PASEF) in a Bruker timsTOF-PRO. (A) Venn diagrams representation of unique and shared proteins found within SG- (blue) and MG-EVs (red). (B) Enrichment analysis of protein classes shared between *A. americanum* SG- and MG-EVs. Bars represent the fold enrichment with p-value ≤ 0.05 and False Discovery Rate (FDR) ≤ 0.05. (C) Western blot analysis of Heat Shock protein 70 (Hsp70) and Calnexin in midgut (MG) and salivary gland (SG) organs (Org) and extracellular vesicles (EVs). Proteins (50 µg for organs and 15 µg for EVs) were separated by SDS-PAGE and transferred into membranes. Numbers next to the ladder represent the size in kilodaltons (kDa). Hsp70 estimated size = ∼70 kDa. Calnexin estimated size = ∼90 kDa.

### 3.2. Unique and differentially expressed proteins within midgut extracellular vesicles mirror organ functions

Although we identified a core set of proteins that were shared between vesicles from different organs, several of the proteins packed within them were unique to each vesicle type [20 proteins in SG-EVs (Supplementary file S3) and 574 in MG-EVs (Supplementary file S4; Figure 1A)]. Nevertheless, the unique proteins in MG-EVs appear to reflect the function of the tissue secreting them. Enrichment analysis showed that the biological processes with the highest fold enrichment in MG-EVs were exocytic process (GO:0140029), vacuolar acidification (GO: 0007035), mitochondrial fusion (GO:0008053), IMP metabolic process (GO:0046040), and clathrin-dependent endocytosis (GO:0072583; Figure 2). Among the enriched molecular functions (Figure S2A), non-membrane spanning protein kinase activity (GO: 0004715), syntaxin binding (GO: 0019905), and sterol transporter activity (GO:0015248) were the most enriched; however, proteins with hydro-lyase activity (GO:0016836) and oxidoreductase activity (GO:0016616) were also enriched. This enrichment suggests a potential role of tick EVs in digestion and detoxification of the blood meal. The enriched protein classes found within MG-EVs are presented in Figure S2B and include heterotrimeric G-protein (PC00117), membrane trafficking regulatory protein (PC00151), dehydrogenase (PC00092), and lyase (PC00144).

**Figure 2.**
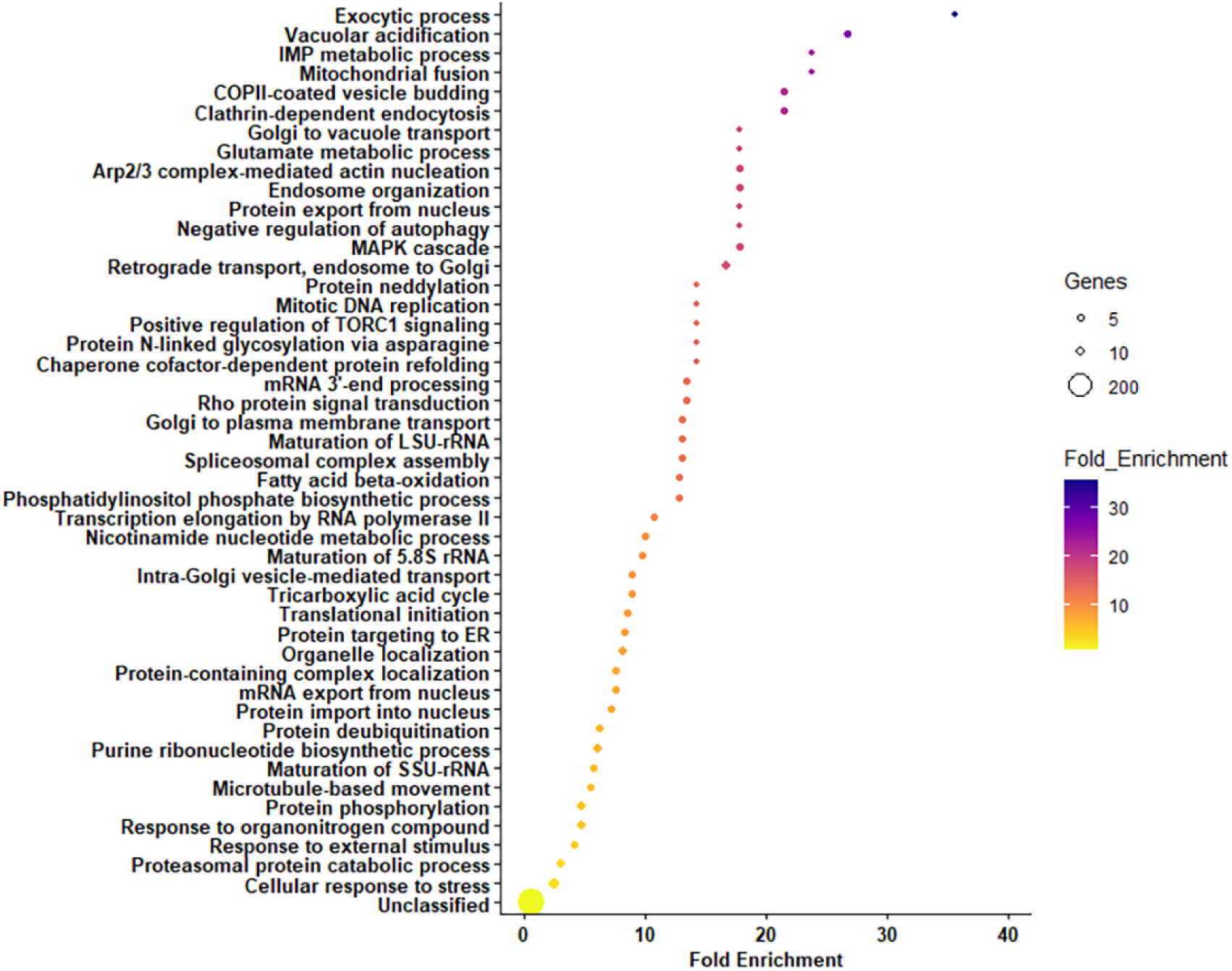
Enrichment analysis of proteins unique to extracellular vesicles (EVs) secreted by midguts from *Amblyomma americanum*. Proteins identified in EVs secreted by *ex vivo* midgut organ cultures from *A. americanum* females fed on white-tailed deer were compared to those from salivary gland EVs to identify unique proteins. An enrichment analysis of biological functions was performed in these unique proteins using Panther. Bars represent the fold enrichment with p-value ≤ 0.05 and False Discovery Rate (FDR) ≤ 0.05.

Of the 1539 proteins detected in both vesicle types, only 516 were quantifiable and 333 were differentially abundant by Student’s T-test q-value (Benjamini Hochberg correction = 0.05). This included 165 proteins more abundant in MG-EVs and 166 proteins were more abundant in SG-EVs (Figure 3A and Supplemental file S5). Enrichment analysis of the proteins differentially expressed in both vesicle populations shows potentially different roles for each vesicle type. MG-EVs were enriched in proteins involved in glutamate metabolic process (GO:0006536), clathrin-dependent endocytosis (GO:0072583), protein N-linked glycosylation (GO:0006487), and other processes (Figure 3B). These processes differed from those proteins over-represented in SG-EVs, which included nucleosome assembly (GO: 0006334), regulation of canonical Wnt signaling pathway (GO:0060828), purine ribonucleoside monophosphate metabolic process (GO:0009167) and others (Figure 3C). Interestingly, except for translation elongation (GO:0006414) and protein folding (GO:0006457), all biological processes enriched within the differentially abundant proteins differed between EV populations, likely due to their distinct function. Likewise, little overlap was observed in Gene Ontology (GO) on the molecular function of proteins differentially abundant in MG-EVs versus SG-EVs, with only actin filament binding (GO:0051015) and translation initiation factor activity (GO:0003743) being shared (Figure S3A and B). Nevertheless, several proteins classes were enriched within the overabundant proteins in both vesicle populations, including translation initiation factors (PC00222), vesicle coat protein (PC00235), RNA splicing factor (PC00148), and RNA helicase (PC00148) (Figure S3 C and D).

**Figure 3.**
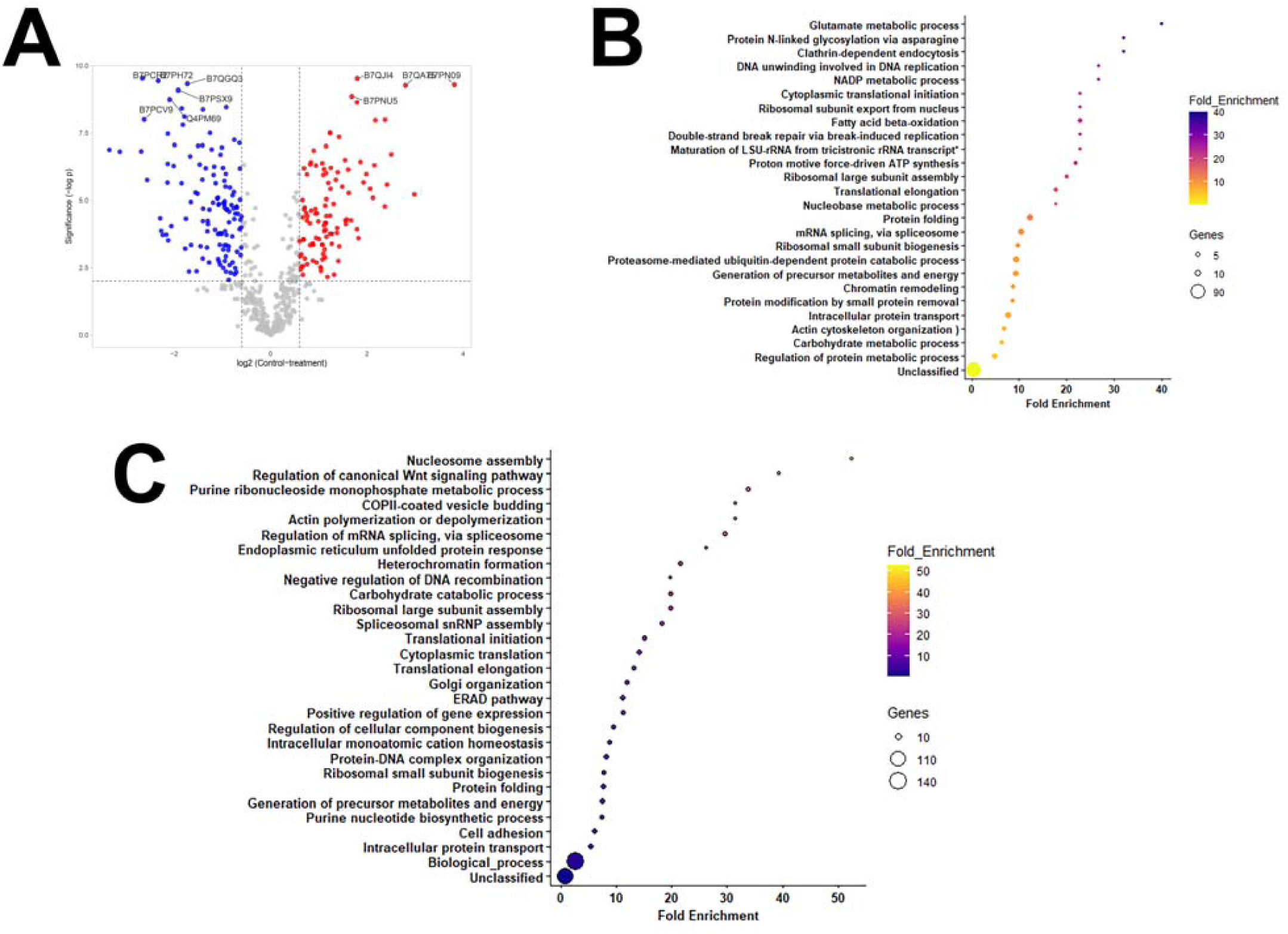
Differentially abundant proteins in extracellular vesicles secreted by midgut and salivary glands from *Amblyomma americanum*. A) Volcano plot of differentially abundant proteins with 1.5-fold cut off. Proteins in blue represent overabundant proteins in salivary glands extracellular vesicles (SG-EVs), whereas proteins in red are more abundant in midgut extracellular vesicles (MG-EVs). Enrichment analysis of the biological function of proteins overabundant in B) MG-EVs or C) SG-EVs. Bars represent the fold enrichment with p-value ≤ 0.05 and False Discovery Rate (FDR) ≤ 0.05.

### 3.3. Salivary gland-derived extracellular vesicles contain more antigenic proteins than midgut vesicles

In our previous study, we visualized several antigenic proteins within MG- and SG-EVs using western blot analysis. This included several bands that appeared consistently in all blots probed with serum from vaccinated animals and serum from animals after tick infestation [22]. To identify these proteins, immuno-precipitations were performed at the Gehrke Proteomics Center at the University of Missouri using pre- and post-vaccination sera from vaccinated WTD #929 and one control WTD (#930), originally. Pre-vaccinated serum was used to identify sticky proteins and proteins naturally recognized by the host. The pre-cleared lysates were then used to precipitate antigenic proteins using the serum of the vaccinated animal (#929) from 7 days after the last booster [22]. The same approach was done using serum from a control animal injected with adjuvant only (WTD #930). These proteins were identified by Data-Dependent Acquisition (DDA) (Supplementary file S6). No proteins were consistently precipitated with the post-injection sera from the control animal (WTD# 930). A total of ten proteins were uniquely found with the post-vaccination serum of WTD #929, including six from SG-EVs and four from MG-EVs. The proteins precipitated from SG-EVs lysates consisted of two alpha-macroglobulins (JAT97670.1 and JAT93800.1), a hypothetical protein (AOE35522.1) that is homolog to a nucleoside diphosphate kinase from *I. scapularis* (XP_002416191.1), a conserved secreted protein (JAT93667.1) without homology to proteins with known function, vitellogenin-1 homolog (AXP34687.1), and an alpha-glucosidase-like enzyme (XP_029834972.1). Antigenic proteins identified from MG-EVs included vitellogenin-2 (JAU01362.1), an acid phosphatase (JAT99430.1), a conserved secreted protein (JAU00577.1), and a hypothetical protein (JAP80714.1) of unknown function.

Due to the drawbacks of the DDA approach, including lower sensitivity, accuracy, and reproducibility [32], the WTD #929 immuno-precipitations were re-acquired using Data-Independent Acquisition (DIA). The identification of antigens precipitated with the serum from vaccinated WTD #924 and #934 and control #930 were also completed using this approach (Supplementary file S7 and S8). DIA allows for unbiased identification of proteins by separating precursors in small windows and identifying all product ions from those windows [32]. Using this approach, we categorized antigens into tree groups: “high-priority”, “medium-priority”, and “low-priority” antigens. Proteins were considered *high-priority* antigens (Table 2) when they were precipitated with post-vaccination serum from treated animals only and had little variation in the spectral intensities, presenting statistical significance. Due to the high variations in the MS-DIA performed with WTD #929, only proteins recognized by WTD#924 and WTD #934 were evaluated in this group (Supplemental file S7). Curiously, only one midgut EV protein was identified, a hypothetical protein (JAT95206.1) with homology to a mucin protein (mucin-2) in *I. scapularis* (XP_029828425.2). In comparison, seven high priority proteins were associated with salivary EVs precipitates, including three enzymes (JAG92630.1, JAT95172.1, and JAU00888.1), two hypothetical proteins (JAT95331.1 and JAP80714.1), none of which have homology with proteins of known function in *I. scapularis*, a conserved secreted protein (JAU00577.1) with no homology to proteins with known function in *I. scapularis*, and protein in meprin (JAU00890.1) (Table 2), homolog to an apical endosomal protein that may function as a low density lipoprotein receptor in *I. scapularis* (XP_040066431.1). These proteins represent strong candidates for future testing.

**Table 1.** Antigenic proteins identified in salivary and midgut extracellular vesicles from *Amblyomma americanum* females by Data-Dependent Acquisition (DDA) using serum from white-tailed deer #929.

| Protein accession # | Protein name (*tick species*) | Vesicle type |
| --- | --- | --- |
| JAT97670.1 | Putative alpha-macroglobulin, partial (*Amblyomma aureolatum*) | Salivary |
| JAT93800.1 | Putative alpha-macroglobulin, partial (*Amblyomma aureolatum*) | Salivary |
| AEO35522.1 | Hypothetical protein (*Amblyomma maculatum*) | Salivary |
| JAT93667.1 | Putative conserved secreted protein precursor (*Amblyomma aureolatum*) | Salivary |
| AXP34687.1 | Vitellogenin-1 (*Haemaphysalis flava*) | Salivary |
| XP_029834972.1 | Lysosomal alpha-glucosidase-like, partial (*Ixodes scapularis*) | Salivary |
| JAU01362.1 | Putative vitellogenin-2, partial (*Amblyomma sculptum*) | Midgut |
| JAT99430.1 | Putative lysosomal acid phosphatase, partial (*Amblyomma sculptum*) | Midgut |
| JAU00577.1 | Putative conserved secreted protein precursor, partial (*Amblyomma sculptum*) | Midgut |
| JAP80714.1 | Hypothetical protein (*Rhipicephalus appendiculatus*) | Midgut |

**Table 2.** High-priority antigens identified in extracellular vesicles secreted by midguts and salivary glands from *Amblyomma americanum* females fed on white-tailed deer.

| Protein accession # | Protein name (*tick species*) | Vesicle type |
| --- | --- | --- |
| JAT95206.1 | Hypothetical protein, partial (*Amblyomma aureolatum*) | Midgut |
| JAG92630.1 | Putative peptidase family m2 angiotensin converting enzyme, partial (*Amblyomma aureolatum*) | Salivary |
| JAT95172.1 | Putative catalytically inactive chitinase-like lectin, partial (*Amblyomma aureolatum*) | Salivary |
| JAT95331.1 | Hypothetical protein, partial (*Amblyomma aureolatum*) | Salivary |
| JAU00577.1 | Putative conserved secreted protein precursor, partial (*Amblyomma sculptum*) | Salivary |
| JAU00888.1 | Putative alpha-d-galactosidase melibiase, partial (*Amblyomma sculptum*) | Salivary |
| JAU00890.1 | Protein in meprin, partial (*Amblyomma sculptum*) | Salivary |
| JAP80714.1 | Hypothetical protein (*Rhipicephalus appendiculatus*) | Salivary |

Proteins that were uniquely precipitated with the vaccinated serum of treated animals, but that show high levels of variation in spectral levels and therefore did not achieve statistical significance were grouped into *medium-priority antigens* (Table 3). As in the previous group, most of the proteins were precipitated from SG-EV lysates (5 proteins). Two midgut proteins were identified, including JAT94005.1, which is a putative secreted mucin that has homology to an elastin-like protein in *I. scapularis* (XP_040078974.1), and JAU02543.1, a hypothetical protein with no homology to proteins in *I. scapularis*. Among the SG-EV proteins precipitated, we discovered three enzymes (JAT97490.1, JAP66750.1, and JAP71591.1), a Hemicentin (XP_029824064.1), and a plasma membrane protein (JAU01951.1), which is a homolog to prominin-1-A [a transmembrane glycoprotein] in *I. scapularis* (XP_042142011.1). The last group of antigens consisted of proteins that were ≥5 fold enriched in the precipitates from vaccinated animals and presented statistical significance in their spectral increases (Table 4). Although it was the largest group of proteins (16 total), these were considered *low priority antigens* as peptides from these proteins were detected in some precipitates from the pre-vaccinated serum. These proteins were all identified in the SG-EV precipitates and included eight enzymes (JAP82974.1, JAT95210.1, AEO36445.1, JAT95556.1, JAU00184.1, JAP80906.1, JAT97770.1, and JAT97963.1), two hypothetical proteins (JAU00361.1 and JAP83809.1), two vitellogenins (JAU01916.1 and AJR36491.1), an alpha-macroglobulin (JAT93800.1), a putative tetraspanin (JAG92577.1), a putative secreted protein (JAT92299.1), with not known function, and a putative protein (AEO35082.1), which is homologous to N(4)-(Beta-N-acetylglucosaminyl)-L-asparaginase in *I. scapularis* (XP_029825395.3).

**Table 3.** Medium-priority antigens identified in extracellular vesicles secreted by midguts and salivary glands from *Amblyomma americanum* females fed on white-tailed deer.

| Protein accession # | Protein name (*tick species*) | Vesicle type |
| --- | --- | --- |
| JAT94005.1 | Putative secreted mucin muc17, partial (*Amblyomma aureolatum*) | Midgut |
| JAU02543.1 | Hypothetical protein, partial (*Amblyomma sculptum*) | Midgut |
| JAT97490.1 | N-acylsphingosine amidohydrolase acid ceramidase, partial (*Amblyomma aureolatum*) | Salivary |
| XP_029824064.1 | Hemicentin-2 putative, partial (*Ixodes scapularis*) | Salivary |
| JAP66750.1 | Metalloprotease atp-dependent zinc metalloprotease yme1 protein isoform x1, partial (*Hyalomma excavatum*) | Salivary |
| JAP71591.1 | LOW glucosidase ii catalytic alpha subunit, partial (*Ixodes ricinus*) | Salivary |
| JAU01951.1 | Putative conserved plasma membrane protein, partial (*Amblyomma sculptum*) | Salivary |

**Table 4.** Low-priority antigens identified in extracellular vesicles secreted by salivary glands from *Amblyomma americanum* females fed on white-tailed deer.

| Protein accession # | Protein name (*tick species*) | Vesicle type |
| --- | --- | --- |
| JAT93800.1 | Putative alpha-macroglobulin, partial (*Amblyomma aureolatum*) | Salivary |
| JAP82974.1 | Uncharacterized alpha-mannosidase (*Rhipicephalus appendiculatus*) | Salivary |
| JAT95210.1 | Putative glucosidase ii catalytic alpha subunit, partial (*Amblyomma aureolatum*) | Salivary |
| JAU00361.1 | Hypothetical protein, partial (*Amblyomma sculptum*) | Salivary |
| AEO36445.1 | Endochitinase-like protein (*Amblyomma maculatum*) | Salivary |
| JAP83809.1 | Hypothetical protein (*Rhipicephalus appendiculatus*) | Salivary |
| JAT95556.1 | Putative aminopeptidase (*Amblyomma aureolatum*) | Salivary |
| JAU00184.1 | Putative catalytic domain of lysosomal alpha-mannosidase, partial (*Amblyomma sculptum*) | Salivary |
| JAP80906.1 | Lysosomal acid phosphatase (*Rhipicephalus appendiculatus*) | Salivary |
| JAG92577.1 | Putative tetraspanin (*Amblyomma americanum*) | Salivary |
| JAU01916.1 | Putative vitellogenin-b (*Amblyomma sculptum*) | Salivary |
| JAT97770.1 | Putative leucine aminopeptidase, partial (*Amblyomma aureolatum*) | Salivary |
| JAT97963.1 | Putative alpha-amylase, partial (*Amblyomma aureolatum*) | Salivary |
| AJR36491.1 | Vitellogenin-6 CP3 (*Ixodes ricinus*) | Salivary |
| JAT92299.1 | Putative secreted protein, partial (*Amblyomma aureolatum*) | Salivary |
| AEO35082.1 | Putative protein (*Amblyomma maculatum*) | Salivary |

## 4. Discussion

Different strategies have been developed for the discovery of anti-tick antigens, including reverse vaccinology based on proteomic and genomic data [33], purification of single proteins from organ and tissue extracts [34], *in silico* immunogenicity analysis of individual proteins [35], or by the selection of individual proteins based on experimental validation of their function [36]. However, some of the drawbacks of these approaches are the time it takes to validate single targets, the lack of the functional characterization of the immune system of WTD for the use of reverse vaccinology and *in silico* testing in this system, and little information on protein expression during tick feeding on wildlife hosts. Unlike in model organisms, only a few studies have characterized genetic and cellular traits associated with immune resistance against parasites and microbes in WTD [37–39]. Further, the genetic structure of the Major Histocompatibility Complex II (MHC II) in ruminants differs from other mammals and WTD have high MHC II polymorphism in *DRB* [40], likely to affect how populations respond to antigens within a vaccine. Given that most vaccine design tools available have been tailored towards humans, animal models, or livestock [41–43], different approaches may be necessary to discover antigens that are reliable for the development of anti-tick vaccines targeting WTD.

We used immunoprecipitation combined with LC-MS/MS to identify the antigenic proteins that activated WTD immune responses after vaccination with tick SG- and MG-EVs. This strategy has previously been employed to pinpoint antigens recognized by the immune response of cattle after vaccination with *I. ricinus* midgut and salivary extracts [44]. Unlike our present study, Knorr *et al.* [44] found around 46 proteins (24 salivary and 22 midgut) that were recognized exclusively by the sera of the animals after immunization and not by pre-vaccination control sera (d0). This is likely due to the higher antigenicity of whole organ extracts compared to extracellular vesicles. Interestingly, several of the proteins that they immunoprecipitated with their immune sera included proteins that we detected within the cargo of EVs in the current study (i.e. heat shock proteins, ribosomal proteins, elongation factors, clathrin, tubulin, histones, etc), indicating that these immunomodulatory proteins are likely secreted within tick EVs, in some cases by both organs. Although many of the proteins reported in Knorr *et al.,* [44] were also immunoprecipitated with the post-vaccination serum for all three WTD, in multiple instances these proteins were also precipitated with serum from our control animal (WTD #930). These included glutathione-S-transferase, which has been previously tested as a vaccine candidate against multiple tick species [45,46], and glyceraldehyde-3-phosphate dehydrogenase in SG-EVs. Our previous western blot using pre-vaccination sera indicated that WTD#930 might have been naturally exposed to ticks or other arthropods with shared homologous proteins [22]. Even though the signal was reduced by day 57, it is possible that some of the circulating antibodies were still present at this time point and precipitated these proteins. Therefore, these proteins should not be discarded as potential candidates for future testing in WTD.

As expected, some of the proteins within our priority lists have previously been tested as antigens or reported as potential antigens for anti-tick vaccines. Knorr *et al.,* [44] identified metalloproteases within their salivary gland extract precipitates. Similarly, we detected a metalloprotease (JAP66750.1) within SG-EVs in our medium-priority antigens. Vitellogenins were also precipitated within our SG-EVs. Among these, vitellogenin-6 had previously been detected in the cement cone of *I. scapularis* ticks [47]. Curiously, several of the targets identified herein were enzymes, including a putative alpha-d-galactosidase melibiase that is of interest as it may naturally lead to the hypersensitivity to red meat that has been associated with *A. americanum* bites. A recent study showed that alpha-d-galactosidase might be involved in the production of N-glycans and other sugar modifications in proteins and other molecules that are injected into the host with tick saliva [48]. Sugar modifications, such as alpha-gal, fucosylation, and sialic acids are important for the transmission of tick-borne pathogens and to stimulate the immune response against ticks [49,50]. Interestingly, silencing of this enzyme in ticks has a favorable effect on tick-feeding, possibly by reducing basophil activation [48]. Basophils are known to be important for anti-tick immunity potentially by interacting with IgE and degranulating [51], which can lead to further cellular infiltration and itchiness. In fact, during experimental infestations in our previous study, we lost many ticks due to grooming [22], likely a pruritic effect of tick feeding.

Nevertheless, most of the proteins discovered, independently of the priority group, in this study consisted of hypothetical proteins (10 total), putative secreted proteins, and other proteins without known function, highlighting our still poor understanding of tick biology. Extracellular vesicles are important for tick feeding [24] and previous experiments showed that ovaries can uptake EVs circulating in tick hemolymph [25]. Interestingly, our proteomic analysis of MG-EVs demonstrates that several of the proteins that are unique to this vesicle type are involved in metabolism, vacuolar acidification, and endocytosis. Ticks complete the digestion of the blood meal intracellularly after digestive cells endocytose the blood meal and form organelles called hemosomes. The blood meal breakdown is done by acidic peptidases [52]. We suspect that unique cargo within MG-EV indicates their potential role in tick digestion. Future experiments should evaluate the expression and function of these proteins, as well as evaluate their efficacy as part of a multivalent vaccine. Extracellular vesicles have a role in activating and modulating immune responses by trafficking and exchanging multiple antigens between cells [53]. The antigens described herein might overcome the disadvantages experienced with previous multivalent vaccine tests, such as antigenic competition [54]. However, future studies should be performed in larger populations of WTD to account for genetic diversity, which might affect vaccine efficacy. One limitation of the present study is the small sample set (three vaccinated and one control WTD), which is due to the complexities of working with ungulate hosts. These results provide proof-of concept that studies to identify tick antigens can be conducted in these targeted host animals, leading to the discovery that EVs 1) have shown protection against ticks in the target host and 2) are known to be expressed during feeding in the target host.

## Supplementary Materials

The following supporting information can be downloaded at: www.mdpi.com/xxx/s1, Figure S1: Enrichment analysis of the biological and molecular functions of core proteins within *Amblyomma americanum* extracellular vesicles; Figure S2: Molecular function (A) and protein class (B) enrichment analysis of proteins unique to midgut extracellular vesicles; Figure S3: Molecular function and protein class enrichment analysis of differentially expressed proteins to midgut (A and C) and salivary (B and D) extracellular vesicles; Supplemental file S1: Normalized spectral intensity from LCMS MG SG EV *Amblyomma americanum*; Supplemental file S2: Salivary and midgut EV shared cargo *Amblyomma americanum* WTD; Supplemental file S3: Proteins unique to *Amblyomma americanum* SG EVs WTD; Supplemental file S4: Proteins unique to *Amblyomma americanum* MG EVs WTD; Supplemental file S5: Differentially abundant proteins; Supplemental file S6: Immunoprecipitation experiment_929-930_SG & MG_Proteins identified_DAA; Supplementary file S7: IP-DIA-924_934_930_MG_SG_pre & vacc_proteins identified; Supplementary file S8: IP-DIA-929_MG_SG_pre & vacc_proteins identified.

## Author Contributions

Conceptualization, AOC.; methodology, AOC; formal analysis, AOC and SD.; investigation, AOC, JG, CH, CSR, BLG, KP, SD, and TLJ.; resources, AOC, PO, and TLJ; data curation, AOC and JG.; writing—original draft preparation, AOC.; writing—review and editing, BLG, JG, CSR, TLJ, and PO.; visualization, AOC.; supervision, AOC and TLJ; project administration, AOC and TLJ.; funding acquisition, AOC and TLJ. All authors have read and agreed to the published version of the manuscript.

## Institutional Review Board Statement

Not applicable.

## Informed Consent Statement

Not applicable

## Acknowledgments

We thank Dr. Brian Mooney at Gehrke Proteomics Center at the University of Missouri for the discussions during the design of the study.

