## Supplementary figures and images for "Identification of *Amblyomma americanum* antigens after vaccination with tick extracellular vesicles in white-tailed deer"

### Figure S1

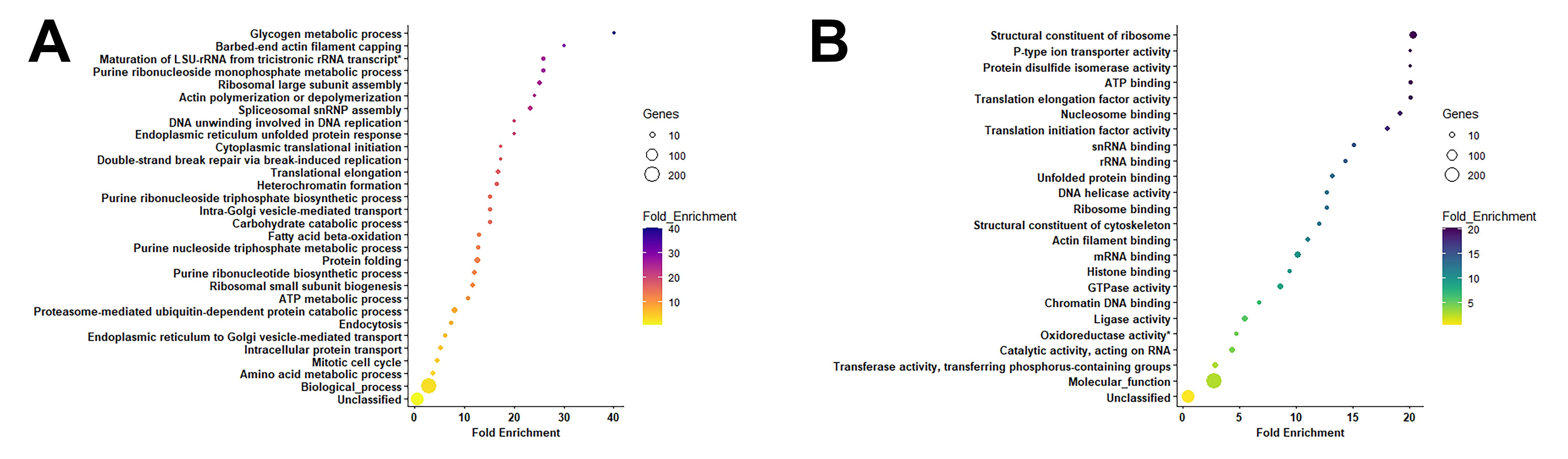

### Figure S2

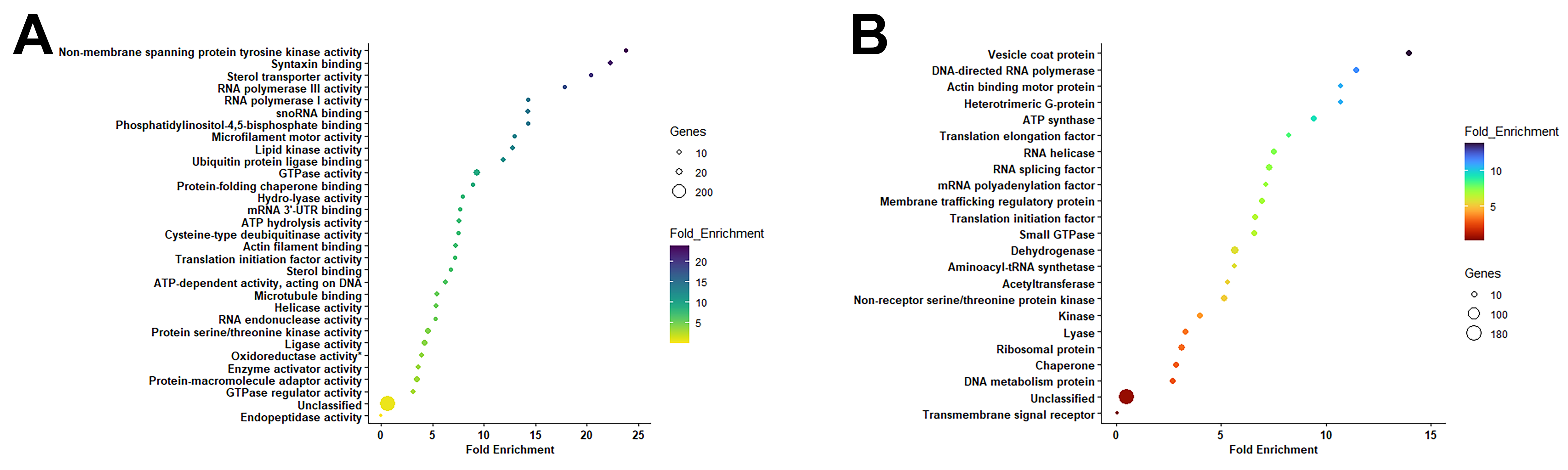

### Figure S3

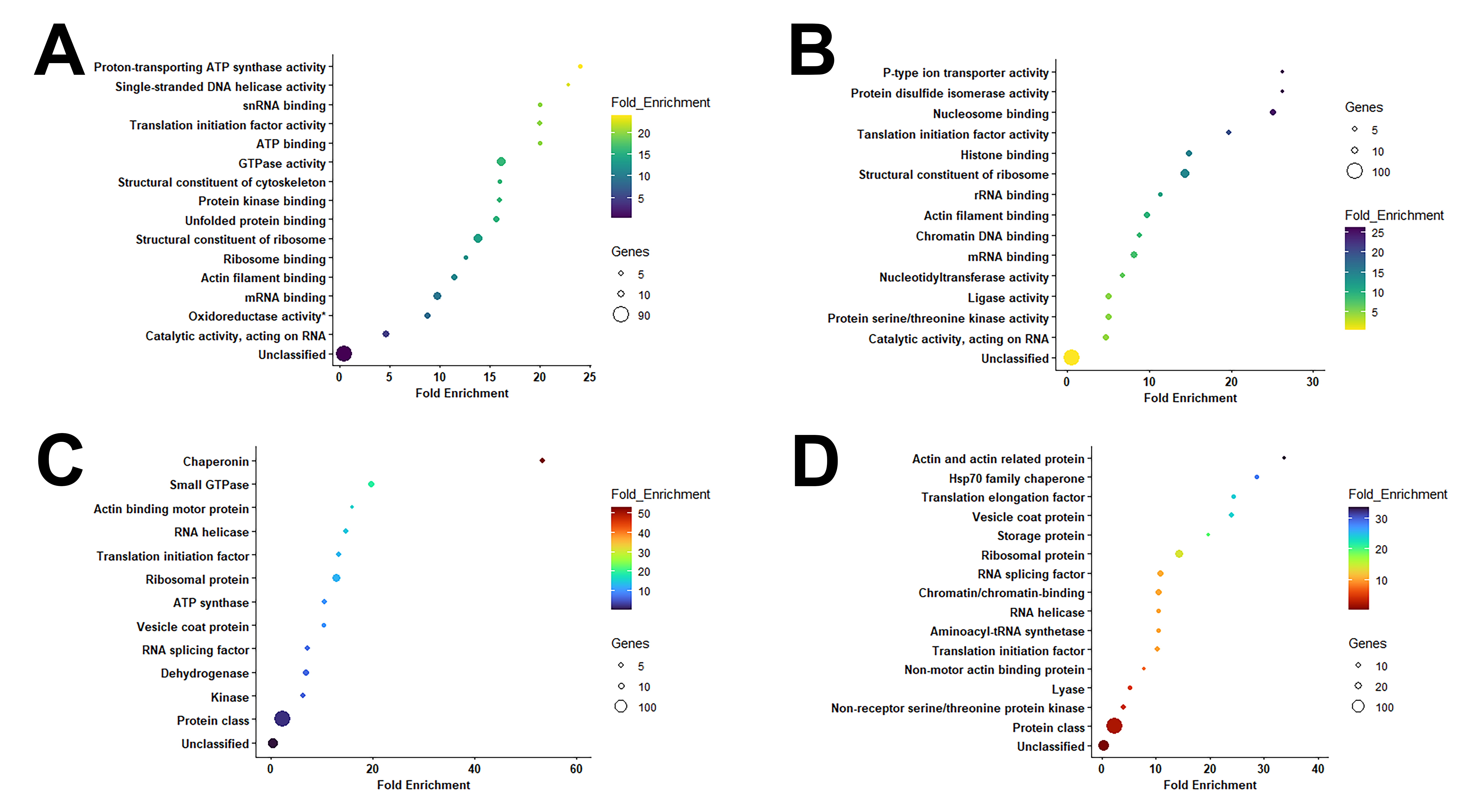
